# A Latent Variable Perspective on EEG Measures of Attentional Control

**DOI:** 10.64898/2026.11.13.766498

**Authors:** Melinda Sabo, Claudia C. von Bastian, Edmund Wascher, Daniel Schneider

**Affiliations:** Leibniz Research Centre for Working Environment and Human Factors, Dortmund, Germany; Yale University, Department of Psychology, USA; University of Sheffield, School of Psychology, UK

**Keywords:** executive functions, cognitive control, P3 component, decoding, structural-equation modeling

## Abstract

Attentional control is a core cognitive function, yet there is still a significant debate revolving around its definition, factorial structure and underlying neural mechanisms. Research in this domain has largely developed along two largely disconnected traditions. Psychometric approaches assess individual differences using latent-variable models (e.g., structural equation modeling, confirmatory factor analysis), while cognitive neuroscience focuses on group-level behavioral and neural effects. This raises a critical question: can neural measures robust at the group level validly capture stable individual differences? Or more specifically, can some of the principles and analytical methods adopted in individual-differences research be directly applied to neural correlates of attentional control? The current study aims to address this question. Electroencephalogram data were recorded from participants as they completed four attentional control paradigms: a change detection task, the go/no-go task, the Simon task, and the Stroop task. From each task, we extracted the P3 component and a data-driven decoding accuracy measure as indices of conflict processing and tested whether these neural measures load onto a common latent factor of attentional control. Results revealed weak and largely non-significant correlations. Structural-equation modeling provided no evidence for a common latent factor, despite robust within-task effects at the group level reflected both in the behavioral and the EEG parameters. These findings suggest that EEG components may lack the construct validity needed to reliably capture individual differences in attentional control.

## Introduction

In our everyday lives, we often find ourselves in situations in which processing certain stimuli in our environment conflicts with our current goals. For instance, while we drive on the highway and search for a specific exit sign, our attention is repeatedly drawn to bright but irrelevant billboards. The ability to regulate information processing in accordance with behavioral goals is commonly referred to as attentional control (also termed cognitive control, executive control, or executive functions; von Bastian et al., 2020). Attentional control can also be conceptualized as a core cognitive function that enables individuals to resolve cognitive conflict by overriding an automatic response in favor of current task demands (Draheim et al., 2021; Oberauer, 2024). This latter definition also highlights how attentional control and conflict processing are intrinsically linked.

Despite its central role in cognition, there is still ongoing debate around the definitions of attentional control and around the models that best describe it. Specifically, substantial disagreement remains about whether it reflects a single underlying ability, multiple distinct but related processes, or both (Oberauer, 2024; von Bastian et al., 2020). Prior work has proposed several competing factorial possibilities. Attentional control may be captured by a single general factor, by multiple correlated first-order factors, by multiple orthogonal factors, or, most radically, may not constitute a coherent psychometric construct at all (Rey-Mermet et al., 2019; von Bastian et al., 2020).

Within this debate, the unity and diversity model has emerged as the most influential framework. Accordingly, attentional control consists of three partially separable but correlated components: shifting, inhibition, and updating. These factors share common variance while also exhibiting component-specific variance (Friedman & Miyake, 2017; Miyake et al., 2000; Miyake & Friedman, 2012). However, recent meta-analytic evidence challenges the robustness of this framework. A large-scale reanalysis of existing datasets found only moderate support for the unity and diversity model, with many studies failing to show clear model acceptance or selection (Karr et al., 2018; Unsworth et al., 2024). These findings suggest that the latent structure of attentional control remains an open question, as to this date no model can be unequivocally accepted (Karr et al., 2018).

A comparable debate revolves around the neural mechanisms underlying attentional control and conflict processing. Converging neural evidence suggests that the fronto-parietal and cingulo-opercular networks orchestrate attentional control and conflict processes (Assem et al., 2020; Freund et al., 2021; C. Gratton et al., 2018; Menon & D’Esposito, 2022). These regions are thought to support core control functions such as conflict detection, conflict monitoring, and regulatory processes (Botvinick et al., 2001) and have led to the proposal of the domain-general account (C. Gratton et al., 2018). At the same time, neuroimaging studies have also provided evidence for task-specific activation patterns within this broader control network, reflecting differences in task demands, stimulus characteristics, or level at which control occurs (Egner et al., 2007; Egner, 2008; C. Gratton et al., 2018; Jiang & Egner, 2014). A third intermediate view posits that neither a fully domain-general nor a fully domain-specific account addresses this question adequately, arguing that the neural implementation of attentional control is characterized by a combination of task-general and task-specific neural modules (Fu et al., 2022; Li et al., 2017; Sabo et al., 2026). Importantly, this view closely parallels the unity and diversity perspective observed at the behavioral level. Indeed, shared and distinct components may coexist not only in performance measures but also in their underlying neural substrates.

Despite substantial behavioral and neural evidence, a critical methodological gap persists in attentional control research (G. Gratton et al., 2018). Individual-differences studies have primarily relied on correlational approaches, in particular latent-variable models (e.g., structural equation modeling, confirmatory factor analysis), to uncover the (latent) structure of attentional control. In contrast, experimental psychology and cognitive neuroscience research has focused on group-level activation patterns. These approaches target the same neurocognitive mechanisms but remain largely disconnected. This raises the question: can neural measures robust at the group level validly capture stable individual differences? Or, more specifically, can some of the principles and analysis methods adopted in individual-differences research be directly applied to the neural correlates of attentional control? The present study aims to bridge this divide by applying latent-variable models from individual-differences research directly to EEG-derived parameters collected across multiple attentional control tasks.

Participants completed four widely used cognitive control tasks: a change detection task (Wascher & Beste, 2010), a go/no-go task, a Simon task (Hoppe et al., 2017; Simon, 1969), and a Stroop task (Stroop, 1935). During task completion, continuous EEG data were recorded, enabling the extraction of multiple neural indices of conflict processing. We focused on two measures. First, we extracted the amplitude of the P3 event-related potential (ERP) component. Although ERP measures such as the N2 and the error-related negativity (ERN) have also been linked to conflict processing in attentional control tasks (Folstein & Van Petten, 2008; Heidlmayr et al., 2020; Meyer & Hajcak, 2019), we did not include them in the current analyses. First, a reliable single-subject ERN estimate requires enough error trials, a condition not met in the Stroop task, where accuracy was at ceiling. Second, given that previous research argues that the Stroop-elicited N2 can be difficult to separate from the later N450 (Folstein & Van Petten, 2008; Liotti et al., 2000; West & Alain, 1999), it is unclear whether the N2-like ERP component captured in the Stroop task indexes the same process as the N2 in other conflict tasks.

Second, we computed decoding accuracy as a data-driven neural index of conflict processing. This measure was obtained by training a linear classifier on raw EEG data to discriminate between conflict and non-conflict trials, yielding an estimate of conflict-related neural processing. Unlike traditional ERP components, this multivariate approach does not depend on predefined time windows or electrode sites and is sensitive to distributed spatiotemporal patterns in the EEG signal. By combining a theoretically motivated ERP component with a data-driven decoding approach, the present study leverages complementary strengths of EEG methodology. This enables us to test whether EEG components of attentional control load onto a common underlying factor and, thus, capture stable individual differences.

## Methods

The data analyzed in the present work were part of the ongoing Dortmund Vital Study (ClinicalTrials.gov identifier: NCT05155397; Gajewski et al., 2022). Within the Dortmund Vital Study, participants completed a vast set of questionnaires and cognitive tasks. Here, we only describe the four tasks included in the analyses. A more comprehensive overview of the full list of cognitive tasks, as well as the data collection procedure can be obtained under: https://vital-study.ifado.de/measures/cognitive-tests-with-eeg-recording. [last accessed on September 24, 2026].

### Participants

Enrollment in the Dortmund Vital Study was restricted to individuals without a history of major medical or psychiatric conditions. Specifically, participants were excluded if they reported neurological disorders (e.g., dementia, Parkinson’s disease, or stroke), cardiovascular disease, bleeding disorders, cancer, or severe psychiatric illnesses such as schizophrenia, obsessive-compulsive disorder, anxiety disorders, or major depression. Additional exclusion criteria included ophthalmological conditions affecting vision (e.g., cataracts, glaucoma, or blindness), prior head trauma, surgical procedures or implants, and limitations in physical fitness or mobility. Individuals currently using psychotropic medication or neuroleptics were also excluded. In contrast, the use of medications such as anticoagulants, hormonal treatments, antihypertensive agents, or lipid-lowering drugs did not preclude participation. All included participants had normal or corrected-to-normal vision and hearing.

Participants’ data were included in the analyses if: (i) they attended both experimental sessions; (ii) they completed all four tasks included in the current study; (iii) their behavioral and EEG recording passed the quality checks; (iv) their vision or hearing impairments did not prevent them from successfully completing the tasks; and (v) no technical issues were encountered during the session. Additionally, participant exclusion was also conducted based on the outlier detection procedure run separately on the behavioral and EEG data for every variable. For the behavioral analyses, the final dataset comprised 278 participants (182 female and 96 male participants, age range: 20-70 years, M_age_ = 40.81, SD_age_ = 12.98), while for the EEG analyses, the data included 305 participants (192 female and 113 male participants, age range: 20-70 years, M_age_ = 43.09, SD_age_ = 13.29).

Prior to study participation, all individuals gave written informed consent. The study was conducted in accordance with the ethical standards set forth in the Declaration of Helsinki and received approval from the ethics committee of the Leibniz Research Centre for Working Environment and Human Factors in Dortmund, Germany (approval no. A93). The tasks were administered across two separate laboratory sessions conducted on two different days. Participants received financial compensation of €160 upon completion of both sessions.

### Experimental procedure

Participants performed a set of computerized cognitive paradigms, comprising a change detection task (Wascher & Beste, 2010), a go/no-go task, a Simon task (Hoppe et al., 2017; Simon, 1969), and a Stroop task (Stroop, 1935; Figure 1). The change detection and Simon tasks were completed during the first experimental session, whereas the Stroop task and the go/no-go task were administered on a separate day. Continuous EEG data were acquired during task completion.

**Figure 1.**
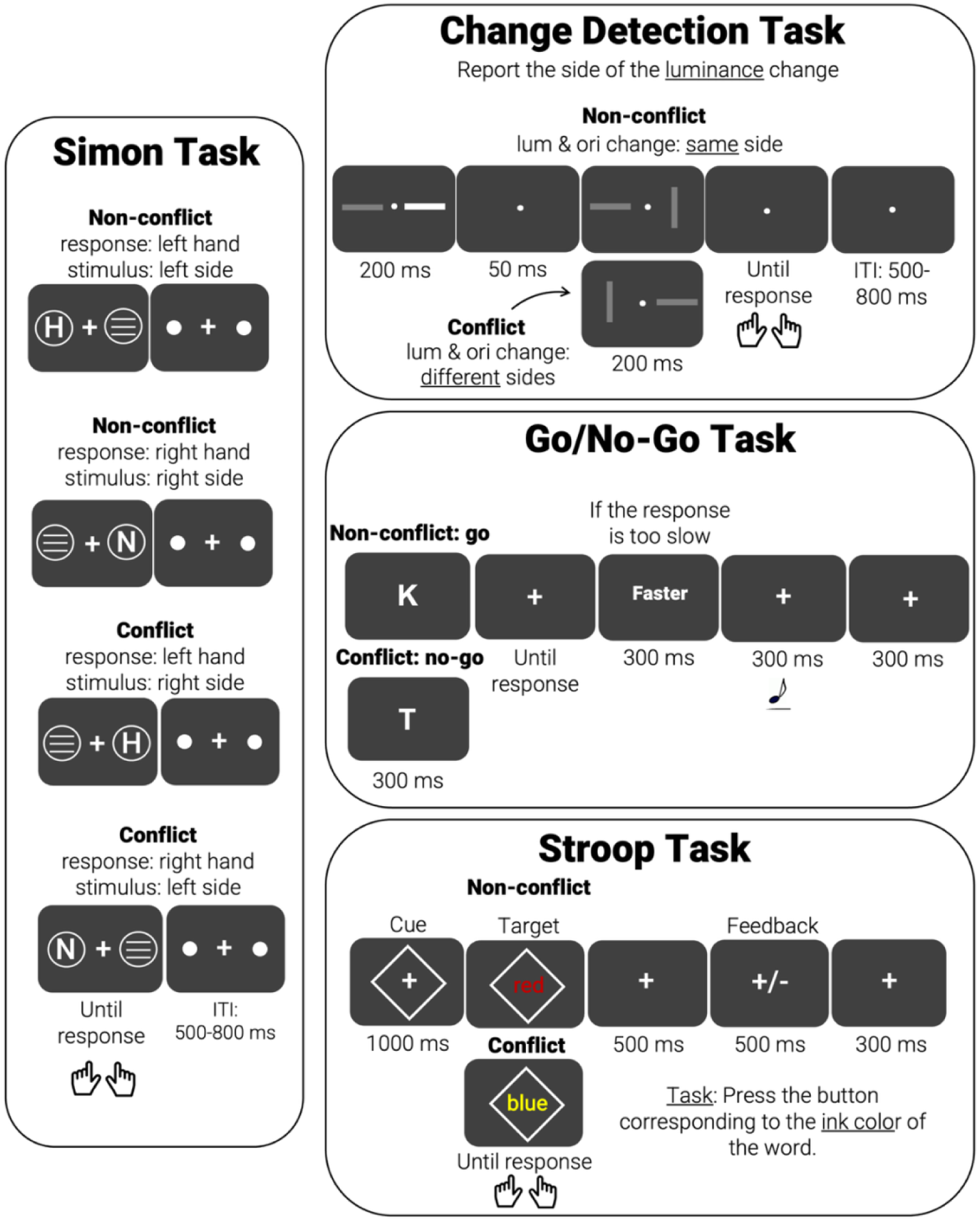
Experimental tasks. <u>Simon task</u>: On each trial, participants first viewed the letter H or N presented on either the left or right side of the screen. They were asked to respond with one hand (e.g., left) to H and with the other hand to N, with hand-letter mappings counterbalanced across participants. Non-conflict trials involved spatially congruent stimulus-response mappings (e.g., left stimulus left-hand response), whereas conflict trials required incongruent responses. <u>Change detection task</u>: Participants were presented with two successive displays, each containing two oriented bars. Their task was to indicate the side on which a luminance change occurred. Some trials also included a task-irrelevant orientation change. Trials consisted of either (a) a luminance change only (non-conflict), (b) luminance and orientation changes on the same side (non-conflict), or (c) luminance and orientation changes on opposite sides (conflict). Responses were made using left- and right-hand buttons corresponding to the side of the luminance change. <u>Go/no-go task</u>: On each trial, participants were shown either the letter K or T, each signaling a “go” or “no-go” instruction. They were instructed to respond as quickly as possible to go-stimuli and to withhold responses to no-go stimuli. Feedback was provided when responses were too slow. Instructions were presented in German. <u>Stroop task</u>: In the color-naming blocks (included in the analyses), participants were asked to report the ink color of the displayed word while ignoring its semantic content. The stimuli consisted of the color words, ‘red’, ‘yellow’, ‘green’, ‘blue’, displayed in German (rot, gelb, grün, blau). The ink and the word’s meaning could be congruent or incongruent. Responses were made using four keys corresponding to the possible ink colors, and feedback was provided after each trial.

**Figure 2.**
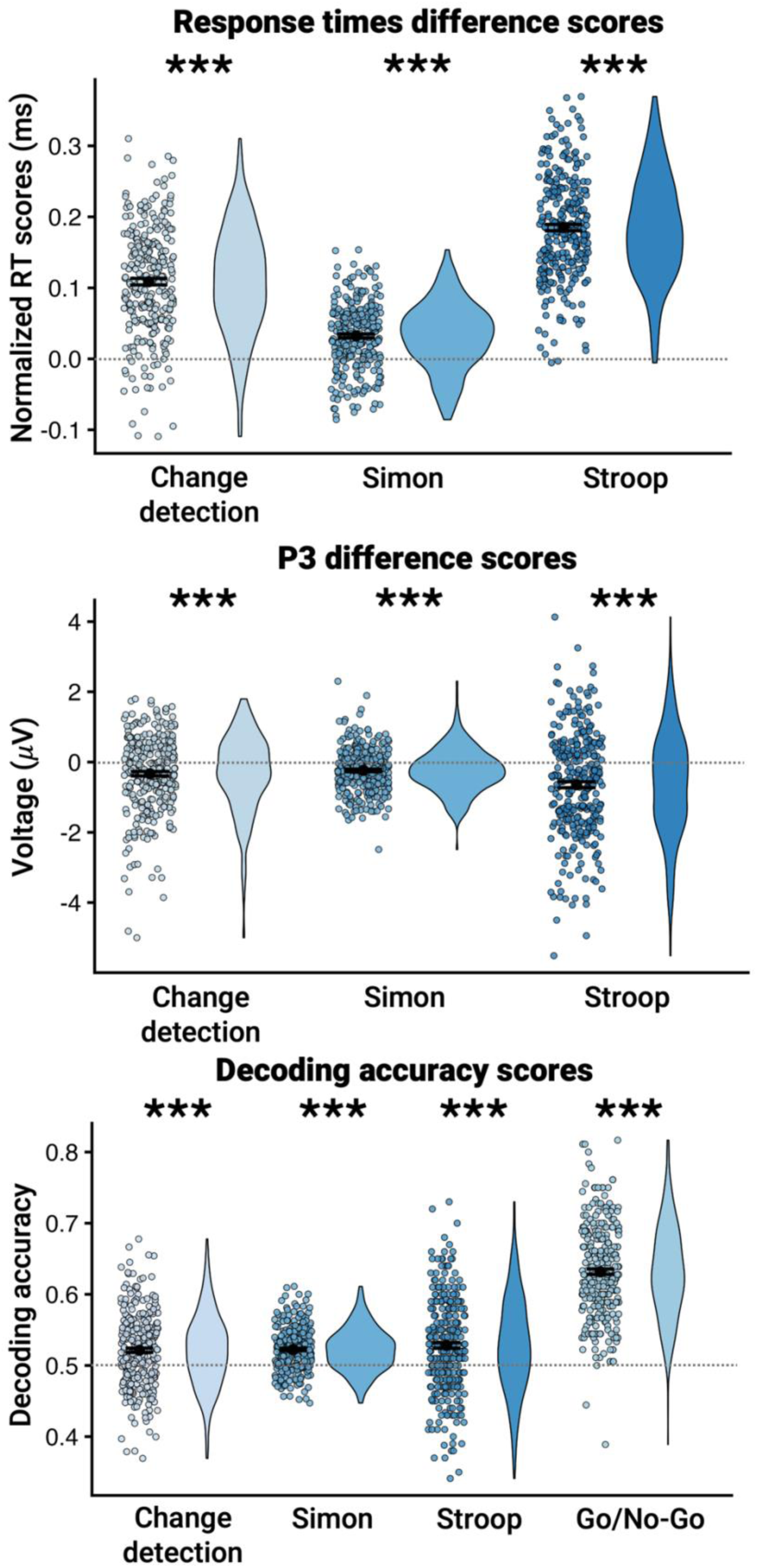
Conflict effects across the three parameters and four tasks. The top panel shows response time difference scores for the change detection, Simon, and Stroop tasks contrasted against zero. The middle panel displays the P3 amplitude difference scores for the same tasks. The bottom panel presents the decoding accuracy scores contrasted against chance level (0.5). Dots represent individual participants, and violin plots illustrate the distribution density of the data. Black horizontal bars indicate group means with corresponding confidence intervals. The dashed horizontal line denotes zero for difference scores (top and middle panels) and chance-level performance (0.5) for decoding accuracy (bottom panel). Asterisks (***) indicate that mean effects significantly differ from zero (or chance level), *p* < .001.

In the first session, stimuli were shown on a 32-inch VSG Display++ LCD monitor (models M0250 & M0251) at a resolution of 1920 × 1080 pixels and a refresh rate of 100 Hz. Manual responses were recorded using force-sensitive response handles, and stimulus generation as well as task control were implemented in FreePascal. EEG data were collected using a 64-channel Ag-AgCl active electrode system (actiCap; Brain Products GmbH), sampled at 1000 Hz and online low-pass filtered at 200 Hz, with FCz used as the reference and AFz as the ground electrode.

In the second session, tasks were displayed on a 17-inch monitor operating at the same refresh rate but a lower resolution of 640 × 480 pixels. Responses were recorded using button-based response devices. EEG recordings were obtained with a 32-channel BioSemi ActiveTwo system (BioSemi B.V.) using Ag-AgCl active electrodes and an online sampling rate of 2048 Hz. The BioSemi system employed a Common Mode Sense active electrode in conjunction with a Driven Right Leg passive electrode to establish the reference and ground via a feedback loop. In case of both EEG sessions, electrodes were positioned according to the international 10–20 system, and impedances were kept below 10 kΩ. On both days, participants were seated at an approximate viewing distance of 70 cm from the screen.

### Simon Task

As shown in the first panel of Figure 1, trials began with a central fixation cross (0.3° × 0.3°) accompanied by two peripheral placeholder dots (0.15° × 0.15°) displayed for 500-800 ms (background: CIE1931: 0.287, 0.312, 10). The subsequent stimulus display presented two shapes, one on each side of a fixation cross. Participants were required to press a button when a circle appeared and withhold responses for diamonds. Only trials requiring a response were analyzed. Circles (1.1° diameter) contained letters on one side, while the opposite shape displayed three horizontal lines (0.45° × 0.07°) as a placeholder (CIE1931: 0.287, 0.312, 80). The letter “H” (0.536° × 0.474°) cued a response with one hand (e.g., right), and “N” (0.495° × 0.474°) cued the other (e.g., left). The letter-hand mapping was counterbalanced across participants. Trials were classified as non-conflict when the target letter appeared on the side corresponding to the correct hand (e.g., left side and left hand) and as conflict when it appeared on the opposite side (e.g., left side and right hand) (Hoppe et al., 2017).

### Change Detection Task

In the change detection task, participants viewed two bars on the left and right side of a fixation cross. After a short delay of 50 ms, another two bars appeared, in which either the luminance, the orientation, or both features of one or both bars were altered. Participants were instructed to report the spatial location (left vs. right) of the luminance change by pressing the corresponding response button. Left side changes were reported with the left hand, while right side changes were reported with the right hand. The visual display’s luminance was set to 20 cd/m². Each bar had a size of 1.35° × 0.56° visual angle and was presented at a distance of 1.3° from the fixation dot (0.3° × 0.3°). The color of the bars was defined in CIE 1931 color space (x = 0.287, y = 0.312), with luminance levels varying between 10 and 50 cd/m². Luminance changes in the second display involved increases or decreases between 10 and 50 cd/m². Orientation changes consisted of transitions between horizontal and vertical alignments (Wascher & Beste, 2010). Feature changes could co-occur within the same bar or be distributed across bars. Based on this manipulation, trials in which the luminance change occurred alone or together with an orientation change in the same bar were classified as non-conflict trials. In conflict trials, the luminance change and orientation change happened at different bars (Figure 1). Importantly, the task also included trials in which there was only an orientation change, which required withholding the response. However, these trials were not included in the current analyses.

### Go/No-Go Task

As depicted in Figure 1, trials started with the presentation of the letter “K” or “T” (size: 3×10 mm). The former indicated that participants had to press the response button (go-trials), while the latter meant that they had to withhold their response (no-go trials). Thirty percent of the trials were no-go trials. If participants did not press the button within 600 ms, they were presented with the written word “faster” (“schneller” in German; size: 5×5 mm), accompanied by a warning sound played for 300 ms. Trials were separated by a 300 ms inter-trial interval during which a fixation dot was displayed (size 5×5 mm).

### Stroop Task

The Stroop task consisted of two primary blocks. In the first block, participants were asked to read the meaning of the presented color words, while in the second block, they reported the ink color of the words. The two blocks were not mixed. The reading block served as the baseline, as it did not involve any conflict, therefore it was excluded from the analyses. Each trial began with a cue, a diamond (3.33° × 3.33°), signaling that the upcoming trial required color naming. After a 1000-ms inter-stimulus interval, a color word was presented (0.57° × 0.82°). Stimuli consisted of the German words “rot,” “grün,” “gelb,” and “blau” (“red,” “green,” “yellow,” and “blue”), each displayed in one of these four ink colors. Trials were either congruent, when the word meaning matched the ink color (non-conflict), or incongruent, when they differed (conflict), with half of the trials in each category. Participants responded using four buttons, each pre-assigned to a specific color, with the index and middle fingers of both hands. All participants used the same color-to-button mapping and had 2500 ms to respond. After response, participants received feedback, during which a plus sign was displayed for 500 ms for correct responses and a minus sign (0.82° × 0.82°) for incorrect responses. This was followed by a 300-ms inter-trial interval (Figure 1). Instructions emphasized both speed and accuracy.

## Data analysish

### Behavioral data

#### Data cleaning

First, we labeled premature and missed responses. In this context, we define premature responses as either button presses made before the relevant stimulus display or those made very shortly after stimulus onset. Premature responses were operationalized as responses preceding a plausible minimum reaction time for each task. A near-zero threshold (−10 ms, allowing for minor stimulus-timing jitter) was chosen for the change detection and Simon tasks. Consistent with established lower bounds for simple visuomotor responses, 80 ms were set for the go/no-go task. Finally, a more conservative 200-ms threshold was selected for the Stroop task, reflecting the additional processing demands of color-word identification and response selection. When it comes to missed responses, we define those as trials in which participants failed to provide a response within a time window. We set 1600 ms for the change detection task, 2900 ms for the go/no-go task, 1500 ms for the Simon task, and 4000 ms for the Stroop task. Premature, missed, and incorrect responses were excluded from the response times analyses. Additionally, we ran an outlier detection procedure on the aggregated data, which removed participants whose values were three median absolute deviations away from the median (Leys et al., 2013). We adopted a conservative approach and ran the outlier detection procedure on the aggregated data for every variable of interest within each task. The outlier detection procedure was run separately on the behavioral and EEG data. The final behavioral data analysis sample comprised 278 participants.

### EEG data

#### Preprocessing

EEG preprocessing was conducted in MATLAB using the EEGLAB toolbox (Delorme & Makeig, 2004). Separate preprocessing pipelines were applied in preparation for the ERP and decoding analyses. In the latter case, EEG preprocessing was deliberately kept to a minimum given the relative insensitivity of multivariate analytical approaches to noise (Carlson et al., 2020). Table 1 provides an overview of the rejected channels and trials.

**Table 1.** Overview of the outcome of ERP preprocessing for each task.

| Task | Number of rejected channels<br>*M* (*SD*) | Percent of remaining trials after trial rejection<br>*M* (*SD*) |
|---|---|---|
| Change detection task | 3.14 (2.57) | 85.19 (6.31) |
| Simon task | 3.96 (2.36) | 87.20 (5.19) |
| Stroop task | 4.77 (2.11) | 95.84 (2.33) |
| Go/no-go task | 3.92 (2.19) | 96.41 (2.07) |

In case of the first pipeline, continuous data were first high-pass filtered at 0.1 Hz and low-pass filtered at 40 Hz. Power line noise and its harmonics were removed using the pop_prepPipeline() function (Bigdely-Shamlo et al., 2015), which also re-references the signal to an estimate of the true average reference and detects and interpolates bad channels. For the independent component analysis (ICA), data were downsampled to 256 Hz, high-pass filtered at 1 Hz, epoched, and baseline-corrected (−200 to 0 ms). Epochs were time-locked to the target stimulus, with task-specific windows: change detection (−500 to 2800 ms), Simon (−500 to 2500 ms), Stroop (−400 to 2000 ms), and go/no-go (−400 to 1200 ms). To ensure that noisy portions of the data do not distort the ICA output, noisy trials were removed via the pop-autorej() function. After the ICA, components were labelled using ICLabel to identify and characterize artifacts, such as eye movement, muscle activity, and heart artifacts (Pion-Tonachini et al., 2019). ICA weights were then applied back to the continuous dataset, and components with ≤30% probability of being brain-related or >30% probability of being eye-related were removed. The cleaned data were finally epoched (same parameters as above), baseline-corrected (−200 to 0 ms), and residual noisy segments were excluded from subsequent analyses.

In preparation for the decoding analyses, continuous data were first band-pass filtered (0.1-100 Hz) using a Hamming-windowed sinc finite impulse response filter and subsequently resampled to 250 Hz. For each experimental paradigm, stimulus-locked epochs were then extracted using the same time windows reported above. Following epoching, baseline correction was applied using the interval -200-0 ms relative to the target item.

#### ERP analysis

Since the main component of interest was the P3, we extracted data from electrode Pz. In the change detection task, the P3 component’s peak was based on the grand average waveform, collapsed across all subjects and conditions, within the predefined time window of 300-600 ms. The peak was defined as the most positive voltage within this window. The component amplitude was then calculated as the mean amplitude in a ±20 ms window centered around the identified peak latency. In the remaining two tasks, we adopted a 50% fractional area latency approach (Liesefeld, 2018; Wascher et al., 2022). This method determines component latency based on the time point at which 50% of the area under the curve is reached within a predefined time window. The following predefined time windows were used: 180-800 ms for the Simon task and 180-1100 ms for the Stroop task. Next, using the cumulative area under the curve, we identified the time point at which 50% of the total positive area was reached. If no clear local maximum was identifiable for an individual dataset, the latency derived from the grand average was substituted. Component amplitude was defined as the mean voltage within ±10 ms around the condition-specific peak identified in the grand average. The peak latencies were identified as follows: 411 ms for the change detection task, 419 ms for the Simon task, 538 ms for the Stroop task. Finally, data were averaged within these windows separately for conflict and non-conflict trials and paired-sample t-tests were used to test for conflict-related EEG effects within each task. Importantly, the go/no-go task was excluded from the P3 analyses, as the go and no-go trials have been argued to elicit different P3 components, with the no-go P3 reflecting a more anteriorly distributed potential (Smith et al., 2008; Verleger et al., 2006).

Decoding analysis. Multivariate pattern classification was carried out in MATLAB using the CoSMoMVPA toolbox (Oosterhof et al., 2016). Across all decoding analyses, classifiers were trained to discriminate between conflict and non-conflict trials, resulting in a chance level of 50% (0.5). A regularized linear discriminant analysis model served as the classification algorithm. To facilitate comparability across participants and recording sessions, datasets were harmonized by restricting analyses to a shared set of electrodes. Training and testing were performed within the same task. Model performance was evaluated using 10-fold cross-validation, in which 90% of trials were used for training and the remaining 10% for testing; this procedure was iterated until each fold had been used as the test set once. Class balance within both training and test partitions was ensured using the cosmo_balance_partitions function, maintaining equal proportions of conflict and non-conflict trials. Because of this step, eight datasets had to be excluded from the decoding analyses, as they did not meet the minimum trial number required for an optimal balancing. Prior to all decoding analyses, a fast Fourier transform was applied to EEG data from a 0–1000 ms post-stimulus window, yielding power estimates across frequencies. Power values in the 1–50 Hz range were extracted and combined with electrode information as features for classification. Feature values were also standardized using z-transformation and these values served as an input for the classifier. Decoding accuracy was quantified as the proportion of correctly classified trials, indexing the classifier’s sensitivity to differences between conflict and non-conflict conditions. The analysis produced a single decoding accuracy score for each participant. To establish that conflict could be successfully decoded within each task, we ran a one-sample t-test relative to chance level.

#### Structural Analyses

The latent structure underlying attentional control was investigated using confirmatory factor analysis within a latent variable framework. All models were estimated in R (version 4.5.2; R Core Team, 2025) using the lavaan package (version 0.6–21; Rosseel, 2012). Model adequacy was assessed based on multiple goodness- of-fit indices, including the chi-square test statistic (χ²), the root mean square error of approximation (RMSEA) with its associated 90% confidence interval, and the standardized root mean square residual (SRMR). Following commonly accepted guidelines, RMSEA values below 0.06 and SRMR values below 0.08 were taken to indicate an acceptable model fit (Hu & Bentler, 1999).

We specified a single-factor model to test whether a common attentional control factor accounted for the observed variance across tasks. We fit this model to normalized response times difference scores (conflict-minus-non-conflict divided by the average of the two conditions), P3 amplitude difference scores (conflict-minus-non-conflict), and decoding accuracy scores separately. We did not fit the model to behavioral accuracy difference scores, because inspection of the Stroop task data revealed that a considerable proportion of participants reached ceiling performance in accuracy (see Figure S1, Supplementary materials). Hence, including this measure with reduced variance would not have allowed for reliable parameter estimation.

In addition, we specified a bifactor model with two orthogonal factors to separate attentional control from general processing speed. Performance across all conditions loaded onto a general processing speed factor, while conflict conditions additionally loaded onto a conflict-specific factor (the go/no-go task was excluded due to its lack of a conflict/non-conflict contrast). Residual errors between conditions of each task were correlated to account for their shared task-specific (method) variance. For decoding accuracy, fitting a bifactor model analogous to the one described above was not feasible. Decoding accuracy is not measured separately for conflict and non-conflict conditions. By construction, the decoding procedure quantifies how well conflict and non-conflict trials can be discriminated from one another based on the EEG signal, so the conflict/non-conflict contrast is already collapsed into a single score per task. Instead, potential confounding effects of response time differences were addressed prior to the analysis by subjecting the signal to frequency decomposition and restricting the analysis to the 0-1000 ms time window, thereby removing the time dimension from the decoding measure itself.

## Results

### Robust Conflict Effects Across All Tasks and Measures

First, we tested the presence of a conflict effect within each task and for each measure (response times, P3 amplitude, and decoding accuracy) included in the modeling procedure. To this end, difference scores between conflict and non-conflict conditions were computed and tested against zero using one-sample t-tests. For the go/no-go task, difference scores could not be derived, as response times were only recorded for go trials. Similarly, no difference scores were computed for the decoding parameter, because decoding accuracy already reflects the classifier’s ability to discriminate between conflict and non-conflict trials. Instead, decoding accuracy was tested against 0.5, representing chance-level performance. For the behavioral measures, analyses focused exclusively on response time differences, as accuracy in the Stroop task was at ceiling (see Figure S1, Supplementary Materials).

Across all tasks and measures, significant effects were observed (see Table 2 for descriptive statistics). For the behavioral measures, participants responded more slowly on conflict than on non-conflict trials, resulting in normalized difference scores that were significantly different from zero: (i) change detection task, *t*(277) = 23.60, *p* < .001, *d =* 1.42, *95% CI* [0.09, 0.11]; (ii) Simon task, *t*(277) = 11.40, *p* < .001, *d =* 0.68, *95%* CI [0.02, 0.03]; (iii) Stroop task, *t*(277) = 41.40, *p* < .001, *d =* 2.48, *95% CI* [0.17, 0.19]. Similarly, P3 amplitudes were more positive for non-conflict than for conflict trials, yielding difference scores that were significantly different from zero: (i) change detection task, *t*(304) = -5.17 *p* < .001, *d =* -0.29, *95% CI* [-0.45, -0.20]; (ii) Simon task, *t*(304) = -6.54, *p* < .001, *d =* -0.37, *95% CI* [-0.30, -0.16]; (iii) Stroop task, *t*(304) = -7.46, *p* < .001, *d =* -0.42, *95% CI* [-0.81, -0.47]. Finally, decoding accuracy was significantly above chance (0.5) for all tasks, indicating that the classifier reliably distinguished between conflict and non-conflict trials: (i) change detection task, *t*(296) = 7.03, *p* < .001, *d =* 0.40, *95% CI* [0.51, 0.52]; (ii) Simon task, *t*(296) = 12.74, *p* < .001, *d =* 0.74, *95% CI* [0.51, 0.52]; (iii) Stroop task, *t*(296) = 7.15, *p* < .001, *d =* 0.41, *95% CI* [0.52, 0.53]; (iv) go/no-go task, *t*(296) = 34.20, *p* < .001, *d =* 1.99, *95% CI* [0.62, 0.63]. Overall, these results confirm that response times, P3 amplitudes, and decoding accuracy all capture reliable differences between conflict and non-conflict trials.

**Table 2.** Descriptive statistics.

| Task | Parameter | M | SD | Min | Max | Skewness | Kurtosis |
|---|---|---|---|---|---|---|---|
| Change detection | normalized RT difference score | 0.10 | 0.07 | -0.10 | 0.31 | -0.17 | -0.08 |
| Go/no-go | RT go-trials | 269.01 | 28.16 | 215.40 | 355.75 | 0.54 | -0.11 |
| Simon | normalized RT difference score | 0.03 | 0.04 | -0.08 | 0.15 | -0.13 | -0.21 |
| Stroop | normalized RT difference score | 0.18 | 0.07 | -0.005 | 0.36 | 0.02 | -0.23 |
| Change detection | P3 difference scores | -0.33 | 1.11 | -5.00 | 1.79 | -0.90 | 1.52 |
| Simon | P3 difference score | -0.23 | 0.62 | -2.48 | 2.30 | 0.20 | 1.04 |
| Stroop | P3 difference score | -0.64 | 1.50 | -5.51 | 4.13 | -0.18 | 0.22 |
| Change detection | Decoding accuracy | 0.52 | 0.05 | 0.36 | 0.67 | 0.08 | 0.20 |
| Go/no-go | Decoding accuracy | 0.63 | 0.06 | 0.38 | 0.81 | 0.03 | 0.20 |
| Simon | Decoding accuracy | 0.52 | 0.03 | 0.44 | 0.61 | 0.24 | -0.03 |
| Stroop | Decoding accuracy | 0.52 | 0.06 | 0.34 | 0.73 | 0.04 | -0.004 |

#### No evidence for a common attentional control factor across tasks and parameters

Table 3 presents the cross-task correlations for each variable of interest. Correlations were generally low across all measures. For the behavioral response time differences, significant associations were observed with age in the change detection, Simon, and go/no-go tasks. In addition, response time differences were significantly correlated between the change detection and go/no-go tasks, *r*(276) = -.15. Correlations among all other tasks were near zero and non-significant. A similar pattern emerged for the P3 component, where correlations between tasks were uniformly small and non-significant. Decoding accuracy correlations were likewise low, with only small positive associations between age and performance in the go/no-go and Simon tasks.

**Table 3.** Cross-task correlations with confidence intervals.

| Variable: normalized RT differences | 1 | 2 | 3 | 4 |
|---|---|---|---|---|
| 1. Age |  |  |  |  |
| 2. Change detection task | -.18**<br>[-.29, -.07] |  |  |  |
| 3. Go/no-go task | .31**<br>[.20, .41] | -.15*<br>[-.26, -.03] |  |  |
| 4. Simon task | .17**<br>[.06, .29] | .01<br>[-.11, .13] | -.03<br>[-.15, .08] |  |
| 5. Stroop task | -.05<br>[-.17, .07] | .01<br>[-.11, .13] | .03<br>[-.08, .15] | .02<br>[-.10, .13] |
| Variable: P3 amplitude differences | 1 | 2 | 3 |  |
| 1. Age |  |  |  |  |
| 2. Change detection task | .01<br>[-.11, .12] |  |  |  |
| 3. Simon task | -.07<br>[-.18, .04] | .04<br>[-.07, .16] |  |  |
| 4. Stroop task | -.08<br>[-.19, .04] | -.06<br>[-.17, .06] | .08<br>[-.03, .19] |  |
| Variable: decoding accuracy | 1 | 2 | 3 | 4 |
| 1. Age |  |  |  |  |
| 2. Change detection task | -.05<br>[-.16, .06] |  |  |  |
| 3. Go/no-go task | .14*<br>[.03, .25] | .02<br>[-.09, .14] |  |  |
| 4. Simon task | .15**<br>[.04, .26] | .05<br>[-.06, .17] | .04<br>[-.08, .15] |  |
| 5. Stroop task | .03<br>[-.08, .14] | .09<br>[-.02, .20] | .10<br>[-.01, .21] | -.05<br>[-.17, .06] |

Note. Values in square brackets indicate the 95% confidence interval for each correlation. \* indicates *p* < .05. \*\* indicates *p* < .01.

Table 4 summarizes all model specifications, modifications, and fit indices. The initial baseline model, which specified a single attentional control factor to capture shared variance across tasks, produced inadmissible solutions. We observed either a negative latent variance for the attentional control factor (response-time difference scores and decoding-accuracy scores) or a negative observed variance for the change-detection task (P3-difference scores). We therefore tested a series of follow-up models to evaluate whether alternative specifications (Models 2 and 3) could yield a viable structure. None of these models produced acceptable solutions, with modifications either resulting in additional Heywood cases, non-convergence, poor model fit, or non-significant factor variances or loadings. Bifactor models (Models 4-6) showed the same pattern: they either failed to converge or produced negative residual variances even when residual variances were constrained to be strictly positive. This suggests that no admissible bifactor solution exists for these data. Together with the low inter-task correlations, these results consistently show that the variance in any of the measures cannot be accounted for by a common latent attentional control factor. Therefore, we conclude that there is no evidence for a domain-general conflict processing factor across our behavioral and EEG measures.

**Table 4.** Overview of tested models.

| Label | Latent Factor(s) | Indicators | Modification | χ² | *df* | *p* | RMSEA | 90% CI | SRMR | Description |
|---|---|---|---|---|---|---|---|---|---|---|
| *Behavioral response times* |  |  |  |  |  |  |  |  |  |  |
| Model 1 | AC | CDT, GNG, SIM, STR | Baseline single-factor model | 0.11 | 2 | .945 | < .001 | [.000, .018] | .006 | Negative AC LV |
| Model 2 | AC | CDT, GNG, SIM, STR | Fix LV to 1 | 0.72 | 3 | .868 | < .001 | [.000, .052] | .016 | Negative CDT OV |
| Model 3 | AC | CDT, GNG, SIM, STR | Fix CDT OV to 1 | 54.89 | 4 | < .001 | .214 | [.166, .266] | .322 | No significant loadings on AC except GNG; poor model fit |
| Model 4 | AC, PS | CDT NC/C, SIM NC/C, STR NC/C | Baseline bifactor model with NC and C conditions loading on PS, and C conditions loading on orthogonal AC, and correlated residual errors between NC/C for each task | 1.95 | 3 | .583 | < .001 | [.000, .086] | .011 | Negative SIM OV for NC and C trials |
| Model 5 | AC, PS | CDT NC/C, SIM NC/C, STR NC/C | Constrain SIM OV for NC and C trials to be positive | 3.98 | 3 | .263 | .034 | [.000, 112] | .035 | Negative residual covariance between SIM OV NC and C trials |
| Model 6 | AC, PS | CDT NC/C, SIM NC/C, STR NC/C | Constrain the residual covariance for SIM NC and C trials to be zero | 4.32 | 4 | .364 | .017 | [.000, .094] | .037 | No significant loadings on and variance of AC LV |
| *EEG P3* |  |  |  |  |  |  |  |  |  |  |
| Model 1 | AC | CDT, SIM, STR | Baseline single-factor model | Model failed to converge |  |  |  |  |  |  |
| Model 2 | AC | CDT, SIM, STR | Fix LV to 1 | 2.28 | 1 | .131 | .065 | [.000, .181] | .035 | Negative CDT OV |
| Model 3 | AC | CDT, SIM, STR | Fix CDT OV to 1 | 61.92 | 2 | < .001 | .313 | [.249, .383] | .413 | No significant loadings on AC; poor model fit |
| Model 4 | AC, PS | CDT NC/C, SIM NC/C, STR NC/C | Baseline bifactor model with NC and C conditions loading on PS, and C conditions loading on orthogonal AC, and correlated residual errors between NC/C for each task | Model fails to converge |  |  |  |  |  |  |
| Model 5 | AC, PS | CDT NC/C, SIM NC/C, STR NC/C | Constrain SIM OV for NC and C trials to be positive | 3.38 | 3 | .336 | .020 | [.000, .101] | .008 | Not positive definite residual covariance matrix due to very low variance in SIM C trials |
| *EEG decoding accuracy* |  |  |  |  |  |  |  |  |  |  |
| Model 1 | AC | CDT, GNG, SIM, STR | Baseline single-factor model | 1.16 | 2 | .560 | < .001 | [.000, .098] | .021 | Negative AC LV |
| Model 2 | AC | CDT, GNG, SIM, STR | Fix LV to 1 | 4.51 | 3 | .212 | .041 | [.000, .114] | .038 | Negative CDT OV |
| Model 3 | AC | CDT, GNG, SIM, STR | Fix CDT OV to 1 | 62.29 | 4 | < .001 | .222 | [.175, .272] | .323 | No significant loadings on AC except STR; poor model fit |

*Note.* AC = attentional control; PS = processing speed; CDT = change-detection task; GNG = go/no-go task; SIM = Simon task; STR = Stroop task; NC = non-conflict; C = conflict; *df* = degrees of freedom; RMSEA = root mean square error of approximation; SRMR = standardized root mean square residual; LV = latent variance; OV = observed variance.

## Discussion

The present study had two related goals. First, we aimed to test whether EEG correlates of attentional control, shown to be robust at the group level, validly capture stable individual differences. Second, we sought to bridge largely separate psychometric and experimental approaches by applying latent variable modeling to established neural indices, testing whether their variance reflects a common underlying factor. To this end, we used the data recorded as part of the Dortmund Vital Study, in which participants completed four attentional control paradigms (change detection, go/no-go, Simon, and Stroop) while their EEG activity was recorded. We integrated behavioral indices (response time difference scores), a canonical ERP component (P3 amplitude), and a data-driven neural metric (decoding accuracy scores), to evaluate whether performance and neural variability across tasks can be accounted for by a single latent attentional control construct.

### Key findings

Across all analyses, we consistently found no evidence for the existence of a domain-general latent factor of attentional control. This was first supported by the correlation analyses, which revealed weak and largely non-significant associations across tasks for both behavioral and neural measures. Furthermore, structural-equation modeling also failed to identify a common latent factor, and this was consistent across response times, P3 amplitudes, and decoding accuracy measures as well as across single-factor and bifactor models.

Our findings are in line with work that does not find a coherent psychometric factor of attentional control (Hedge et al., 2018; Paap & Sawi, 2016; Rey-Mermet et al., 2018, 2019). In addition, the current study extends recent efforts to test whether electrophysiological indices of attentional control can load onto a common factor of attentional control. For example, recent work examining the N2 and P3 components across multiple inhibition tasks (arrow flanker, negative priming, and Stroop) found that the P3 did not reliably differentiate conflict from non-conflict trials at the group level (Löffler et al., 2025). This work suggested that inhibition-related P3 variance was fully accounted for by a general, task-unrelated factor. Our findings extend these results. Despite robust conflict vs. non-conflict differences in the P3 amplitude and successful decoding of the conflict signal within each task, these measures showed no evidence of a shared latent factor across tasks.

The absence of evidence for a unitary model in the neural parameters may be partly attributable to conflict arising at different processing stages across tasks. In the Simon task, there is broad consensus that conflict primarily emerges later, at the response-selection stage, reflecting competition between the stimulus location and the response hand (e.g., Leuthold, 2011). Similarly, in the go/no-go task, conflict is typically linked to response-related processes, particularly the need to inhibit prepotent responses. In contrast, the locus of conflict in the Stroop task remains more debated, with accounts arguing for both stimulus- and response-level conflicts (Doehrman et al., 1978; Hock & Egeth, 1970; MacLeod, 1991; Steinhauser & Hübner, 2009; Van Veen & Carter, 2005; Zysset et al., 2001). A similar multi-stage interpretation may apply to the change detection task. On the one hand, participants must resolve a perceptual conflict between task-relevant and task-irrelevant feature changes. On the other hand, because responses are mapped onto different hands, the task also entails response selection between competing alternatives, introducing a Simon-like response conflict. The four tasks can thus be ordered by conflict locus, from predominantly response-related (Simon, go/no-go) through mixed (change detection, Stroop).

This account makes a testable prediction. Specifically, it suggests that tasks sharing a similar conflict locus should correlate more strongly with each other than with tasks that do not. Our results revealed a significant RT correlation between the change detection and go/no-go tasks, which both entail response-related conflict. However, the correlation between the Simon and the go/no-go tasks or the Simon and the change detection tasks did not reach significance. This suggests that processing stage may shape task-specific neural signatures, but it does not fully explain the lack of shared variance.

Another potential factor contributing to the lack of shared variance across tasks concerns the inherent complexity of trial-averaged conflict estimates. Trial-to-trial sequential dependencies, such as congruency-sequence (conflict adaptation) effects, are a well-documented feature of conflict paradigms (e.g., Braem et al., 2014) and have been shown to shape the reliability and validity of the resulting attentional control measures (Moretti et al., 2026). Averaged conflict and non-conflict estimates may therefore reflect a blend of conflict-detection and conflict-adaptation-related processes whose relative contribution is difficult to isolate and may vary across paradigms, potentially obscuring shared, task-general variance.

#### Implications of the current study

The current results have broader implications for individual-differences research, particularly with respect to the use of neural markers of attentional control. Specifically, our results imply that EEG components examined in the present study may lack the construct validity required to serve as reliable measures of individual differences, even though the experimental effects are robust and consistent for each task. A critical issue is that experimental, group-level and individual-differences approaches are sensitive to fundamentally different aspects of the data (Friedman & Miyake, 2017). Group-level analyses of neural data are designed to detect effects that are consistent across participants. However, precisely because such neural signals are similar across individuals, they may have little explanatory power for individual differences in behavior. In contrast, neural processes that do contribute to variability in performance are, by definition, those that differ between individuals. Such processes may exhibit substantial inter-individual variability and, consequently, may fail to reach statistical significance in traditional group-level analyses, where variability is treated as noise. This asymmetry has important consequences. It implies that the neural signals most prominently identified in experimental cognitive neuroscience are not necessarily those that are most informative for understanding stable individual differences in attentional control. Conversely, signals that are potentially important for explaining such differences may remain undetected when relying solely on group-level contrasts.

This tension echoes a long-standing divide within psychology. Cronbach (1957) described experimental and individual-differences research as two traditions that developed largely in isolation, famously noting that the methodological gap between them left "the country between” these two "turn into desert" (Cronbach, 1957, p. 673) that needs traversing. Bridging this divide has remained a persistent challenge. Recent work has raised this concern explicitly for attentional control, cautioning that paradigms optimized to isolate robust, replicable group-level effects are not automatically well suited to capturing stable individual differences (von Bastian et al., 2020). Our findings extend this argument to the neural level. EEG measures of conflict processing appear to face the same fundamental tension that Cronbach identified for behavioral measures. Bridging this gap will require approaches that explicitly integrate these perspectives, for instance by prioritizing measures that capture both within-task reliability and between-task variability, or by developing models that more directly link task-specific neural processes to stable individual differences.

## Conclusion

Using behavioral and EEG data recorded in four attentional control tasks, we investigated whether common EEG parameters of attentional control can be characterized as a unitary psychometric construct. Correlation analyses revealed weak and largely non-significant associations across tasks and structural equation modeling consistently failed to identify a common latent factor across all behavioral and neural parameters. Together, these findings suggest that EEG components may lack the construct validity required to serve as a reliable measure of individual differences, even though at the group level and within each task the effects are robust and consistent. Moreover, these results may challenge the assumption that different EEG correlates of attentional control capture a unitary, domain-general neural mechanism.

## Acknowledgments

The authors would like to thank the Dortmund Vital Study team for their support with conducting telephone interviews, organizing and conducting the testing and data preprocessing: Ludger Blanke, Tobias Blanke, Claudia Brockhaus, Pia Deltenre, Barbara Foschi, Silke Joiko, Karin Lukaszewski, Carola Reiffen, and Christiane Westedt. The study was endorsed by the German Center for Mental Health (DZPG).

## Conflict of interest

The authors declare no conflicts of interest.

## Use of artificial intelligence

During the preparation of this manuscript, the authors used (Claude, Opus 5.5, Anthropic) to refine the language and improve the readability of the text. The same tool was also used to generate the R script for the behavioral figure (Figure 2). The authors reviewed, edited and checked all AI-assisted content, including the code and the figure it produced. The authors take full responsibility for the content of this publication.

## Notes

### Competing Interest Statement

The authors have declared no competing interest.

