## Supplementary materials for "A Latent Variable Perspective on EEG Measures of Attentional Control"

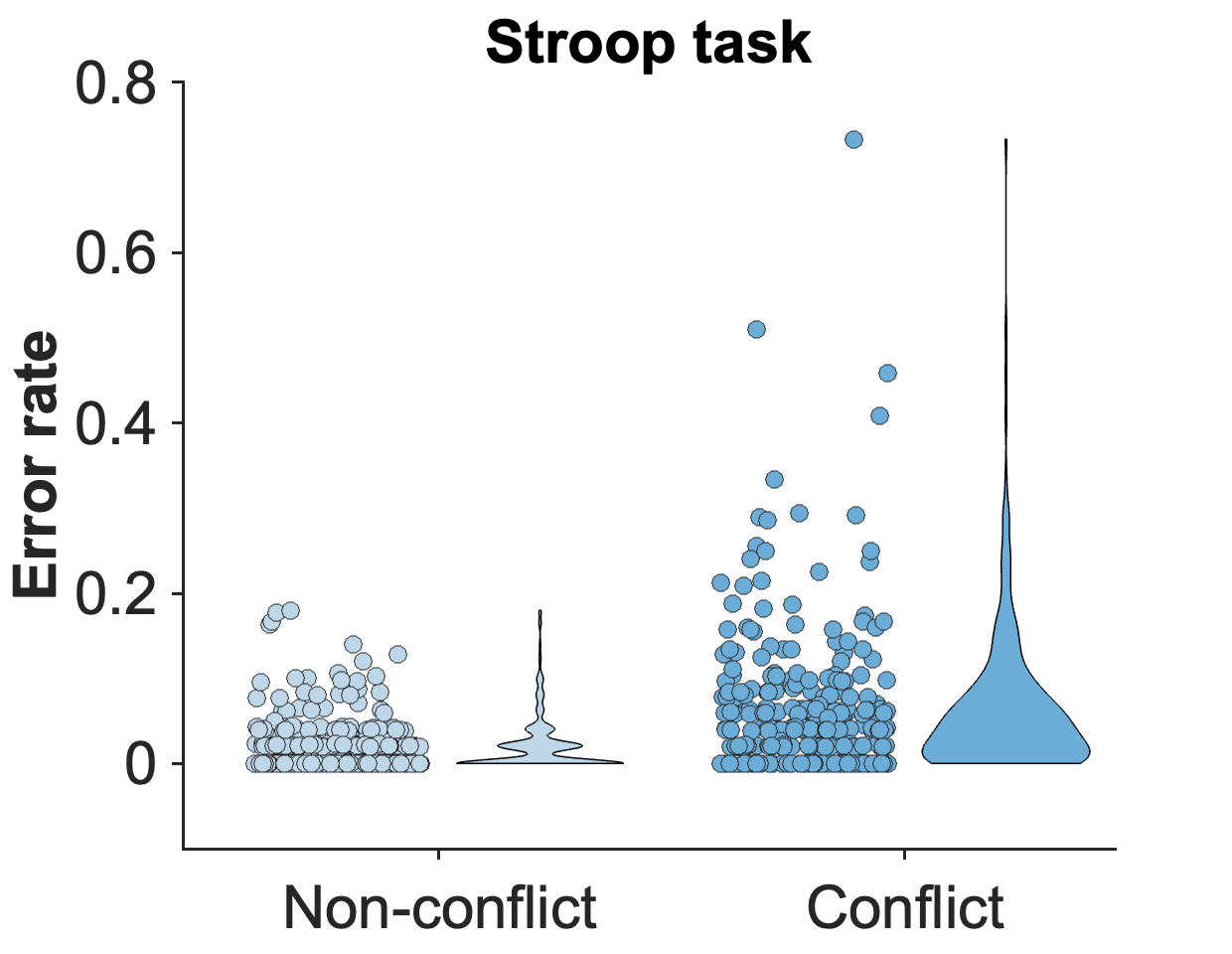
Melinda Sabo ^1,2^, Claudia C. von Bastian ^3^, Edmund Wascher ^1*^, Daniel Schneider ^1*^

**Figure S1.** Error distribution in the Stroop task. Dots represent individual participants, and violin plots illustrate the distribution density of the data. Participants’ performance is at ceiling with a considerable proportion of participants being at an error rate of 0.
